# Designing novel biochemical pathways to commodity chemicals using ReactPRED and RetroPath2.0

**DOI:** 10.1101/2020.12.31.425007

**Authors:** Eleanor Vigrass, M. Ahsanul Islam

## Abstract

Commodity chemicals are high-demand chemicals, used by chemical industries to synthesise cocountless chemical products of daily use. For many of these chemicals, the main production process uses petroleum-based feedstocks. Concerns over these limited resources and their associated environmental problems, as well as mounting global pressure to reduce CO_2_ emissions have motivated efforts to find biochemical pathways capable of producing these chemicals. Advances in metabolic engineering have led to the development of technologies capable of designing novel biochemical pathways to commodity chemicals. Computational software tools, ReactPRED and RetroPath2.0 were utilised to design 49 novel pathways to produce benzene, phenol, and 1,2-propanediol — all industrially important chemicals with limited biochemical knowledge. A pragmatic methodology for pathway curation was developed to analyse thousands and millions of pathways that were generated using the software. This method utilises publicly accessible biological databases, including MetaNetX, PubChem, and MetaCyc to analyse the generated outputs and assign EC numbers to the predicted reactions. The workflow described here for pathway generation and curation can be used to develop novel biochemical pathways to commodity chemicals from numerous starting compounds.

## Introduction

Commodity chemicals, such as ethylene, propylene, benzene, phenol, ethanol, and toluene, are high-value chemicals used by industries to synthesise countless chemical products of daily use. From pharmaceuticals to biofuels (Bengelsdorf and Dürre, 2017; Straathof, 2014), the global chemical turnover was valued at € 3475 billion in 2017, and this demand is expected to rise further in the future (Cefic, 2018). Both organic and inorganic commodities are mainly derived from fossil fuel-based petroleum feedstocks to release harmful direct and indirect greenhouse gases such as CO_2_ and CO into the atmosphere. Concerns over these limited fossil-fuel resources and increasing global pressure to reduce greenhouse gas emissions (UNEP, 2017) have led to an urgent need to find sustainable biochemical routes capable of producing these chemicals and satisfying their demands.

Biochemical routes involving fermentation and enzymatic methods have widely discussed in the literature for sustainable production of commodity chemicals (Saha, 2003; Siebert and Wendisch, 2015). Fermentation is a microbial process that uses microorganisms such as bacteria and yeast to produce enzymes (Renge et al., 2012), which then catalyse the biochemical reactions producing commodities from sugars and other biomass resources (Straathof, 2014). For example, the production of ethanol via the fermentation of syngas (Bengelsdorf et al., 2013; Bengelsdorf and Dürre, 2017), or the conversion of protein waste to cinnamic acid and β-alanine (Kumar et al., 2015) are microbially mediated fermentation processes. Enzymes are highly selective, but they also have the ability of catalyse numerous non-selective or non-specific reactions in addition to the specific reaction the enzyme has evolved for (Kumar et al., 2015; Straathof, 2014). This ability of catalysing non-specific reactions is known as the ‘enzyme promiscuity’ (Tawfik, 2010), and is dependent on the substrates and cofactors involved in the reactions (Delépine et al., 2018; Shin et al., 2013; Tawfik, 2010). Although billions of years of evolution have enriched the repertoire of natural biochemical reaction networks of an organism, many chemical commodities cannot be produced ‘naturally’ due to surpassing an organism’s natural capabilities (Wang et al., 2017). Additionally, there is lack of knowledge on promiscuous enzyme activities such as the number of promiscuous reactions that enzymes can partake (Lee et al., 2012; Shin et al., 2013; Wang et al., 2017). These limitations prevent the discovery and implementation of potential biochemical pathways to high-value commodity chemicals.

Recent advances in cheminformatics and bioinformatics have enabled the design of novel (i.e., biologically unknown) biochemical pathways (Brunk et al., 2012; Medema et al., 2012), and have expanded our knowledge of promiscuous enzyme activities through the design and implementation of computational tools (Hadadi et al., 2019; Wang et al., 2017). Many of these state-of-the-art computational tools are equipped with unique abilities to aid metabolic engineering efforts by designing novel pathways for numerous applications, including bioremediation of xenobiotics (Finley et al., 2009), novel drug discovery (Moura et al., 2016), and production of commodity chemicals (Islam et al., 2017; Yim et al., 2011). Examples of some of the widely used cheminformatics tools include From Metabolite to Metabolite (FMM) (http://fmm.mbc.nctu.edu.tw/), BINCE (Hatzimanikatis et al., 2005), DESHARKY (Rodrigo et al., 2008), PathPred (Moriya et al., 2010), and MRE (Kuwahara et al., 2016). These tools have been applied to numerous studies and have been extensively discussed elsewhere (Brunk et al., 2012; Henry et al., 2010; Islam et al., 2017; Medema et al., 2012; Wang et al., 2017).

Many of these computational tools ‘retrosynthetically’ generate biochemical pathways by iteratively applying the ‘generalised reaction rules’ to transform and connect target compounds to the metabolites of interest (Hadadi et al., 2016; Medema et al., 2012; Wang et al., 2017). The generalised reaction rules are derived using the EC (Enzyme commission) number information of known biochemical reactions assigned by the Nomenclature Committee of the International Union of Biochemistry and Molecular Biology (NC-IUBMB, 1992). These tools have the capability of generating novel and known biochemical reactions; however, a significant limitation that most tools suffer from is the combinational explosion of pathways predicted due to using the generalised reaction rules. The number of pathways generated could result in the thousands and in some cases, in millions, presenting the challenge of efficient post-processing of the generated pathways to find meaningful results (Islam et al., 2017). Although publications relevant to a specific software provide information on how to use and generate results using the software, often there is no further guidance on how to curate these results to obtain useful pathways: a crucial need for practicing metabolic engineers. This need leads to developing individual curation methods that are mainly tools or software specific, as well as specific to the conducted studies.

In this study, two powerful computational cheminformatics tools, ReactPRED (Sivakumar et al., 2016) and RetroPath2.0 (Delépine et al., 2018) were applied to design novel biochemical pathways to produce three commodity chemicals: benzene, phenol, and 1, 2-propanediol. These target compounds were chosen based on their limited biochemical pathway knowledge (i.e., how many pathways are known in the current biological databases) and global demand. For example, it was estimated that the global demand for benzene in 2016 was 46 million tonnes (Pérez-Uresti et al., 2017). RetroPath2.0 and ReactPRED are relatively new, open source, and customisable cheminformatics tools. We chose to use these tools based on their ability to predict novel retrosynthetic (i.e., transforming the target compounds to their simpler precursors) and synthetic (i.e., using simpler precursor compounds to construct target molecules) pathways through identifying the chemical bond transformations occurring in the reactions (Delépine et al., 2018; Sivakumar et al., 2016). After generating pathways automatically, a pathway curation method was developed for each tool to analyse millions of generated pathways and remove redundant results. Initially pathways were examined for specific starting compounds, such as acetate, pyruvate, and glucose, as these compounds are abundantly available in cells and widely used in their metabolisms. Next the pathways containing these compounds were screened based on thermodynamic feasibility. The feasible pathways were further analysed to examine the compounds generated, and individual reactions were assigned to an enzyme commission (EC) number: a numerical classification scheme for enzyme catalysed reactions (Egelhofer et al., 2010). This task was accomplished by comparing the generated reactions to known reactions in the MetaNetX (Moretti et al., 2016), MetaCyc (Caspi et al., 2020) and KEGG (Kanehisa et al., 2020) databases. Finally, both software programmes were analysed to discuss their comparative advantages and limitations for finding novel biochemical pathways to target compounds.

## Materials and methods

### Automated generation of pathways

Novel biochemical pathways were constructed for the production of 1, 2-propanediol, benzene, and phenol using the computational software programmes, ReactPRED and RetroPath2.0. Detailed descriptions of both algorithms and their functionalities can be found elsewhere (Delépine et al., 2018; Sivakumar et al., 2016). Both software programmes require a number of inputs to generate biochemical pathways. In the case of ReactPRED, these inputs included a set of generalised reaction rules developed based on the EC numbers of biochemical reactions found in the MetaCyc (Caspi et al., 2020) database, cofactors (NAD, NADP), target compounds (benzene, phenol, 1, 2-propanediol), and source (glucose, pyruvate, acetate) compounds information in the SMILES format (Weininger, 1988). For RetroPath2.0, the inputs included a set of reaction rules generated based on the biochemical reactions in the MetaNetX (Moretti et al., 2016) database and the reactions in the genome-scale *E. coli* metabolic model, iJ01366 (Orth et al., 2011), as well as the source, sink, and cofactor compounds, including benzene, phenol, NAD, and NADP. RetroPath2.0 then retrosynthetically and ReactPRED synthetically generated pathways by iteratively applying the generalised reaction rules to generate reactions connecting the target compounds to the metabolites present within the MetaCyc and MetaNetX databases. The thermodynamic feasibility of both ReactPRED and Rertopath2.0 generated pathways was analysed by estimating the standard Gibbs free energy of the generated reactions using the group contribution method (Jankowski et al., 2008; Noor et al., 2012).

### Manual curation of the generated pathways

The automatically generated pathways were analysed and manually curated based on the reactions and compounds involved in reactions, and the overall pathway feasibility. For RetroPath 2.0, most of the generated reactions that were biologically known were automatically assigned an EC number. However, the unknown or novel reactions were examined and compared to similar reactions in the MetaNetX (Moretti et al., 2016) database. Also, the generated compounds were all examined to verify their identities by comparing them with the compounds in the MetaNetX and PubChem (Kim et al., 2019) databases. If the compounds were identified and existed in the databases, they were assigned to corresponding reactions with an EC number while unconfirmed compounds were discarded (Figure 1). ReactPRED generated compounds in the SMILES format (Weininger, 1988), which were examined and verified using the PubChem database. Unidentified compounds were discarded while identified compounds were further assessed using the MetaCyc (Caspi et al., 2020) and KEGG (Kanehisa et al., 2020) databases to confirm if the generated compounds were present in the biological databases. The identified compounds and corresponding reactions were then assigned an EC number based on the reaction rule and cofactor information used to generate those reactions. Moreover, the compounds not present in MetaCyc and KEGG were further analysed with the online CDK depicter tool (Willighagen et al., 2017) to confirm the bond transformations occurring within the proposed reactions (Figure 1). An EC number was then assigned to the reactions using the provided reaction rule and cofactor information.

**Figure 1:**
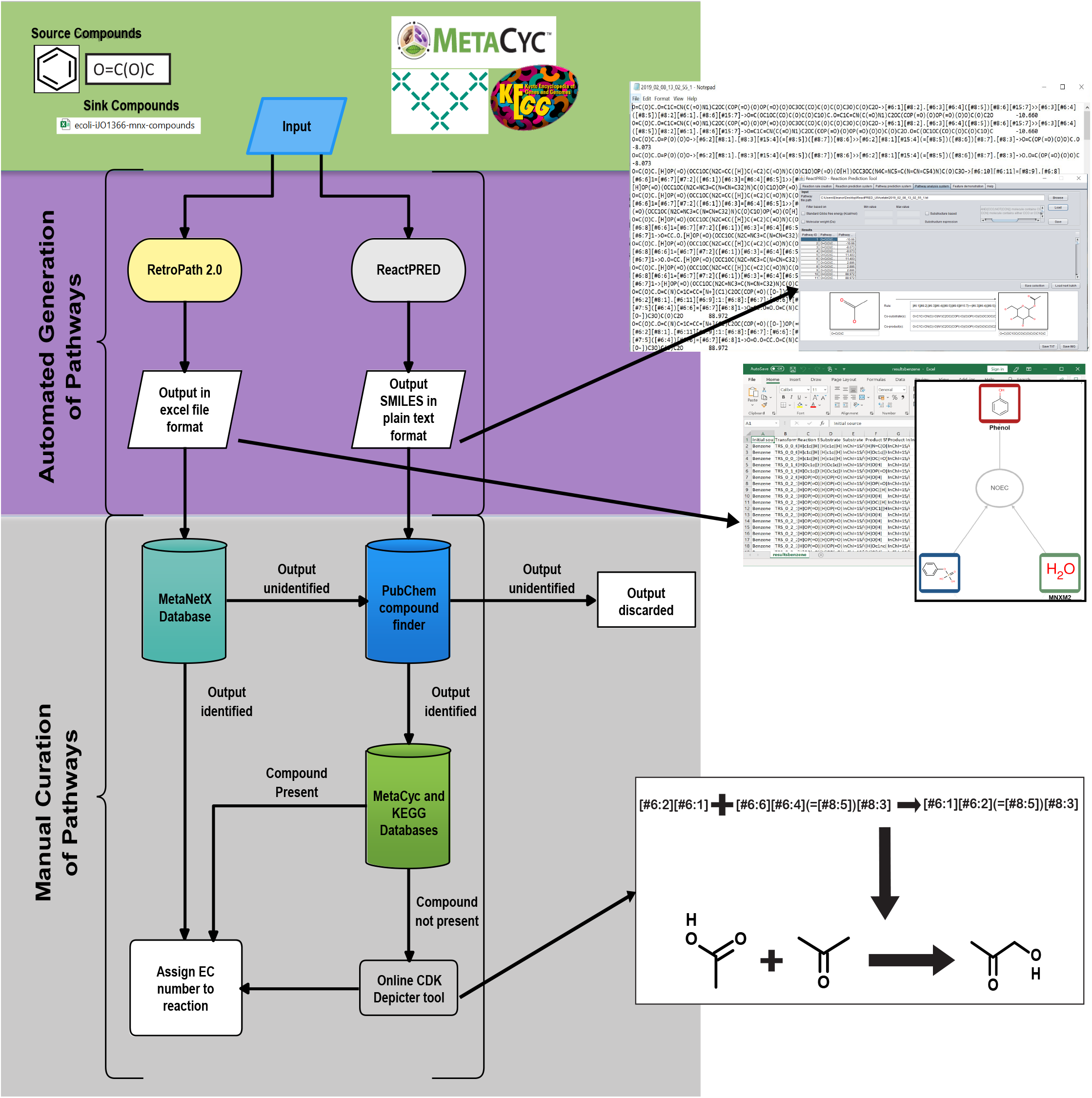
Schematic of the overall workflow followed to design biochemical pathways using RetroPath2.0 and ReactPRED. The input to both software tools included source or target compounds, sink compounds, and reaction rules developed from biochemical databases and genome-scale *E. coli* metabolic model, iJO1366. The generated outputs in excel (RetroPath2.0) and SMILES (ReactPRED) formats were manually curated and extensively analysed with MetaNetX, PubChem, MetaCyc, and KEGG databases, as well as with the online CDK depicter tool to remove non-natural compounds and assign EC numbers to accepted reactions (see text for details).

## Results and discussion

### Analysis of the generated pathways using ReactPRED

Reactions and pathways were generated using ReactPRED’s default reaction rule set, which included a total of 1462 reaction rules (Sivakumar et al., 2016) and the SMILES format of the starting compounds. Glucose, pyruvate, and acetate were used to generate synthetic pathways up to the pathway length of 3, while phenol, benzene, and 1, 2-propanediol were used as starting compounds to retrosynthetically generate pathways up to the pathway length of 2.

Figure 2 illustrates the number of pathways generated with increasing pathway length. The number of generated pathways is linked to the number of potential bond transformations available in the starting compound. For example, the number of glucose pathways produced at each pathway length is greater than the number of acetate and pyruvate pathways produced (Figure 2A). Additionally, the number of phenol pathways significantly increased from 1952 at pathway length 1 to 2337181 at pathway length 3 (Figure 2B), further illustrating that more potential for bond transformations in an input compound generates more outputs. From the generated pathways, thermodynamically feasible pathways, i.e., reactions with a negative standard Gibbs free energy to the target compounds were examined. Table 1 shows the number of thermodynamically feasible pathways generated to the target compounds at each pathway length. No pathways were found to synthetically generate benzene. Table 2 shows the number of thermodynamically feasible retrosynthetic pathways generated at each pathway length.

**Figure 2:**
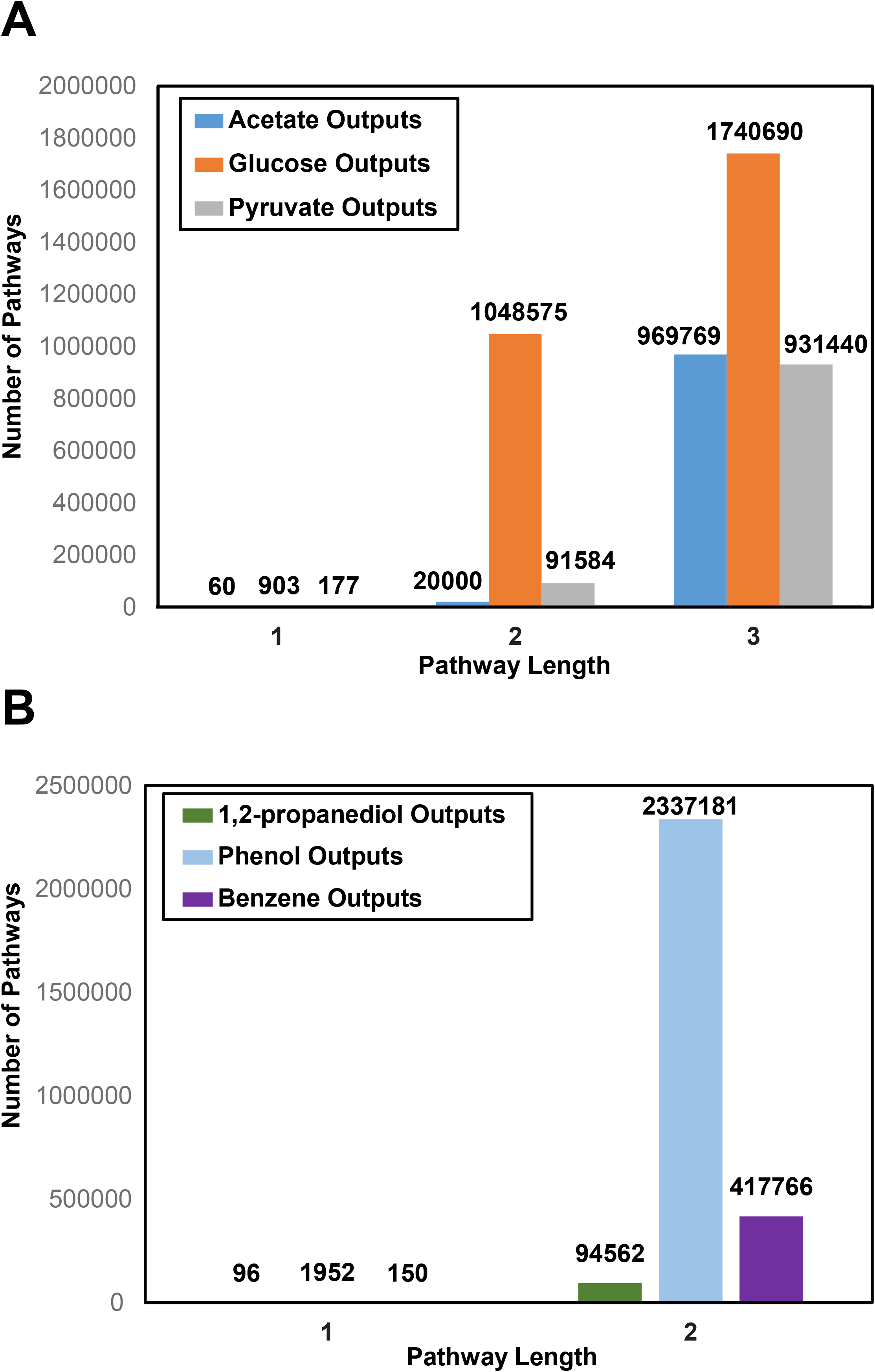
Total number of pathways generated by the ReactPRED software. (A) is showing the number of pathways generated synthetically using acetate, glucose, and pyruvate as starting compounds, and (B) is showing the number of retrosynthetic pathways generated using 1,2-propanediol, phenol, and benzene as starting compounds. The total number of compounds that were generated at each pathway length are also illustrated.

**Table 1:** Number of thermodynamically feasible and synthetically generated pathways to target chemicals (1,2-propanediol, benzene, phenol) from synthetic starting compounds (acetate, pyruvate, glucose)

| Pathway Length | Thermodynamically feasible pathways to 1,2-Propanediol | Thermodynamically feasible pathways to Benzene | Thermodynamically feasible pathways to Phenol |
| --- | --- | --- | --- |
| 1 | 0 | 0 | 1 |
| 2 | 1 | 0 | 21 |
| 3 | 2 | 0 | 164 |

**Table 2:** Number of thermodynamically feasible and retrosynthetically generated pathways to target compounds (acetate, glucose, pyruvate) from retrosynthetic inputs (1,2-propanediol, benzene, phenol)

| Pathway Length | Thermodynamically feasible pathways to Acetate | Thermodynamically feasible pathways to Glucose | Thermodynamically feasible pathways to Pyruvate |
| --- | --- | --- | --- |
| 1 | 0 | 1 | 0 |
| 2 | 2220 | 3580 | 57 |

Comparing both the synthetic and retrosynthetic results, more pathways were generated retrosynthetically because there were more potential for bond transformations in the retrosynthetic starting compounds than the compounds used for the synthetic analysis. Therefore, more reaction rules were automatically applied to these compounds to generate more outputs, i.e., reactions. Pathways were further analysed individually based on the identity of the compounds involved in the pathways (Figure 3). Pathways were discarded if they included compounds unidentifiable in the PubChem database (Kim et al., 2019). For instance, many of generated compounds contained carbon atoms with 5 or more bonds, which means these compounds would be unlikely to exist in nature. From the synthetic outputs, two pathways to 1, 2-propanediol and one pathway to phenol were accepted (Figure 3A). Figure 3B shows the number of accepted and discarded pathways to each target compound for the retrosynthetic outputs: one acetate, ten pyruvate, and seven glucose to 1, 2-propanediol pathways were accepted, while seven acetate, five pyruvate, and fifty nine glucose to phenol pathways were accepted. No acetate to benzene or pyruvate to benzene pathways were accepted because each of these categories of pathways included at least one unidentifiable compound; however, fifteen glucose to benzene pathways were accepted. The accepted reactions were assigned EC numbers capable of catalysing the reactions following the procedure described in Materials and Methods.

**Figure 3:**
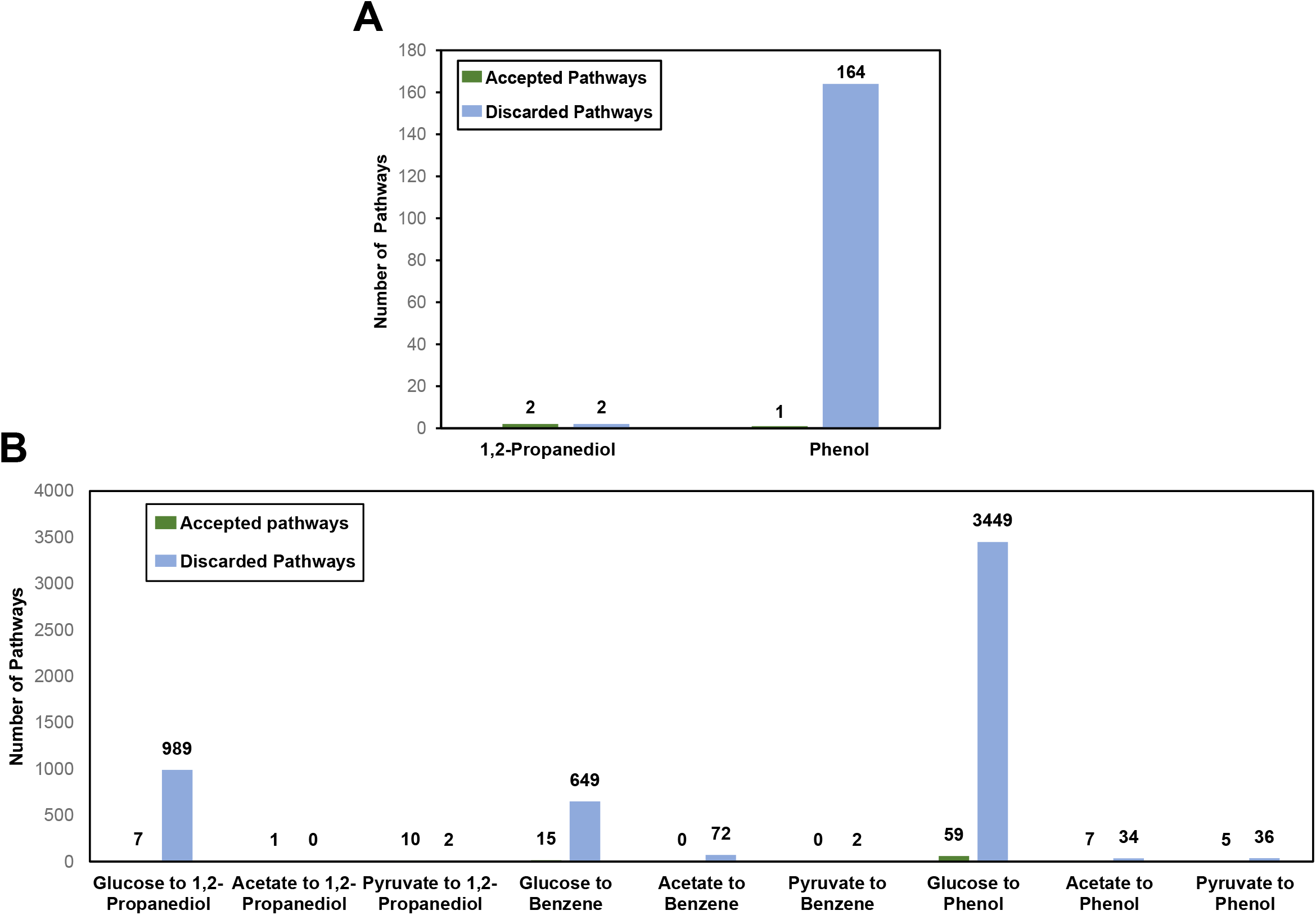
The number of accepted and discarded pathways to the target chemicals obtained from synthetic and retrosynthetic runs of ReactPRED software is shown in (A) and (B), respectively. The thermodynamically feasible pathways were subjected to further pathway pruning (see materials and methods). The accepted pathways are the ones that successfully passed the curation criteria while the discarded pathways failed to pass the curation criteria (see materials and methods).

### Analysis of the generated pathways using RetroPath2.0

The RetroPath2.0 algorithm generated pathways using 14,302 reaction rules known to the *E. coli* metabolism, benzene as the starting compound, and a set of ‘sink’ compounds. The sink compounds are the native metabolites of the chassis organism, i.e., *E coli* metabolism (Delépine et al., 2018). The algorithm converged after 7 iterations, as no new reaction was generated from further run of the algorithm (Figure 4). Figure 4A illustrates the total number of generated reactions, the number of reactions assigned to EC numbers, and the number of compounds generated at each iteration. The number of generated compounds and reactions peaked at iteration 3, generating in total thirty seven reactions and eighty nine compounds, while the total number of reactions and compounds significantly decreased to three and five after the third iteration. This reduction in compounds and reactions can be attributed to the handling of ‘sink’ compounds by the algorithm. Ordinarily, the total number of reactions should exponentially increase with each iteration of the algorithm. However, RetroPath2.0 removes all outputs, in which the generated compounds match those that are in the sink set (Delépine et al., 2018); thus, preventing further iterations of the algorithm on those generated compounds. Each reaction was further analysed, and reactions were discarded if they contained compounds unidentifiable in the PubChem (Kim et al., 2019) database. Thus, no reactions were accepted from those that were generated after the fourth iteration (Figure 4B). However, three, two, and twelve reactions were accepted from the reactions generated after the 1^st^, 2^nd^, and 3^rd^ iteration of the algorithm, respectively (Figure 4B). RetroPath2.0 automatically assigns EC numbers to each reaction. Only one accepted reaction from the first iteration set and ten accepted reactions from the third iteration set were assigned to EC numbers. The unassigned accepted reactions were then compared to the ReactPRED results to identify similar reactions. Additionally, the reaction rule and co-substrate information were examined to manually assign an EC number to the unassigned reactions.

**Figure 4:**
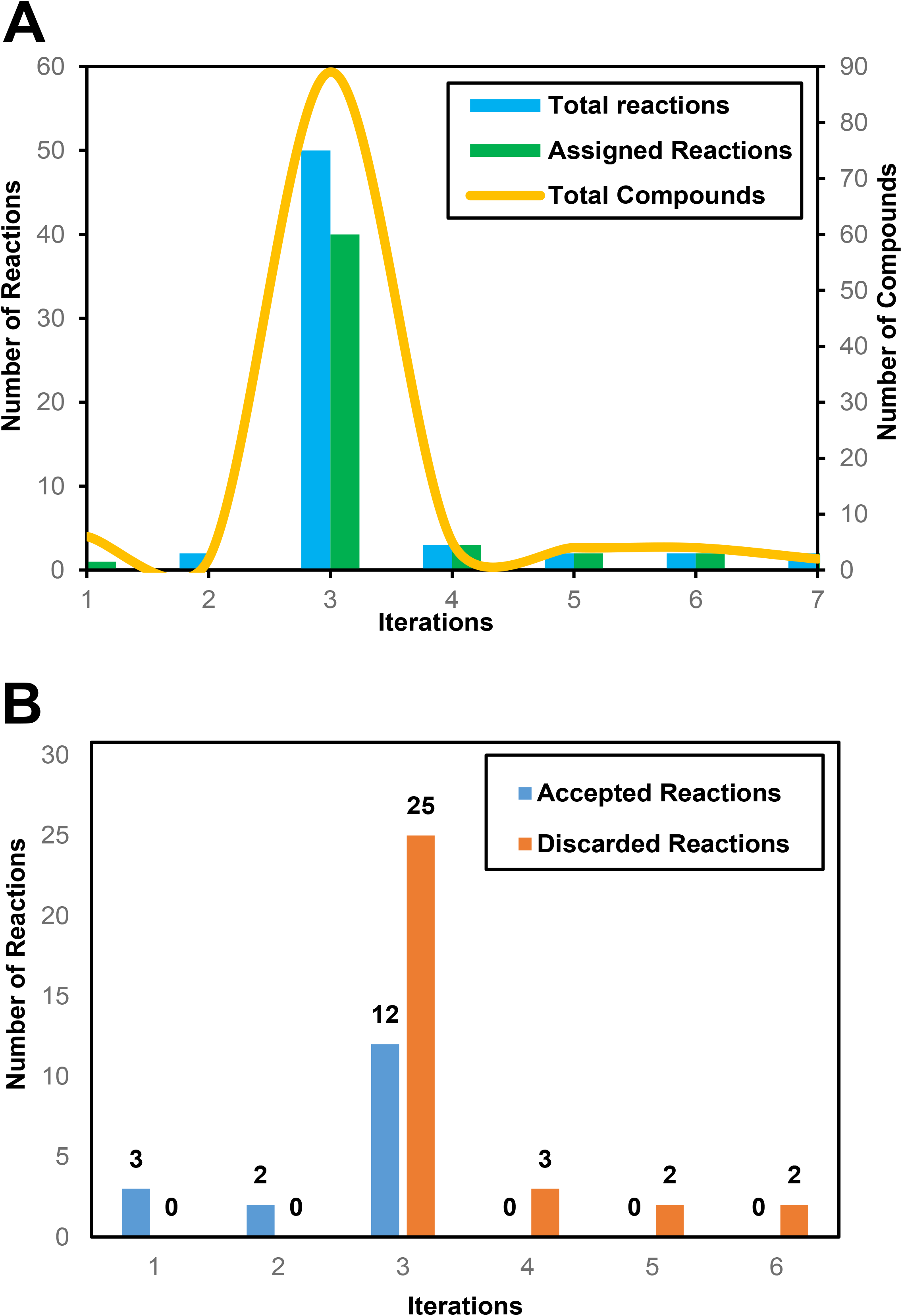
Illustration of the results generated by the RetroPath2.0 software: (A) is showing the number of reactions and compounds generated, as well as EC numbers assigned to generated reactions using this software. Notably, 76% of the total reactions were assigned an EC number at iteration 3 (represented by the green bar). (B) is showing the number of accepted and discarded reactions generated at each iteration of RetroPath2.0. The predicted reactions were subjected to further pruning (see materials and methods). The accepted pathways are the ones that successfully passed the curation criteria, while the discarded pathways failed to pass the curation criteria (see materials and methods).

No direct pathways to glucose, acetate, or pyruvate were found using RetroPath2.0. However, pathways were generated to connect compounds present within the *E. coli* metabolism, as well as in the PubChem database only. Additionally, generating pathways retrosynthetically using phenol as the starting compound produced the same results as benzene, but no pathways were generated using 1, 2-propanediol as the starting compound.

### Analysis of the accepted pathways

From the generated outputs of ReactPRED and RetroPath2.0, in total 49 pathways consisting of 106 reactions connecting acetate, glucose, and pyruvate to phenol, benzene, and 1, 2-propanediol were accepted. No 1-step pathway connecting benzene or 1, 2-propanediol to the target starting compounds, i.e., glucose, acetate, and pyruvate was identified. Each pathway contains at least one novel step, while 25 (51%) of the accepted pathways are entirely composed of novel reactions. No pathways were identified in which all reaction steps were known, i.e., found in the MetaCyc, MetaNetX, and KEGG databases. Many of the accepted pathways contained identical reaction rules, as well as compounds found in the PubChem database only. For example, 13 of the 26 (50%) glucose to phenol pathways (Supplementary data) contained compounds only identifiable in the PubChem database. Many of these compounds are synthetic man-made compounds that are found only in the retrosynthetically generated pathways. Figure 5 shows the number of accepted pathways from glucose, pyruvate, and acetate to each commodity chemical.

**Figure 5:**
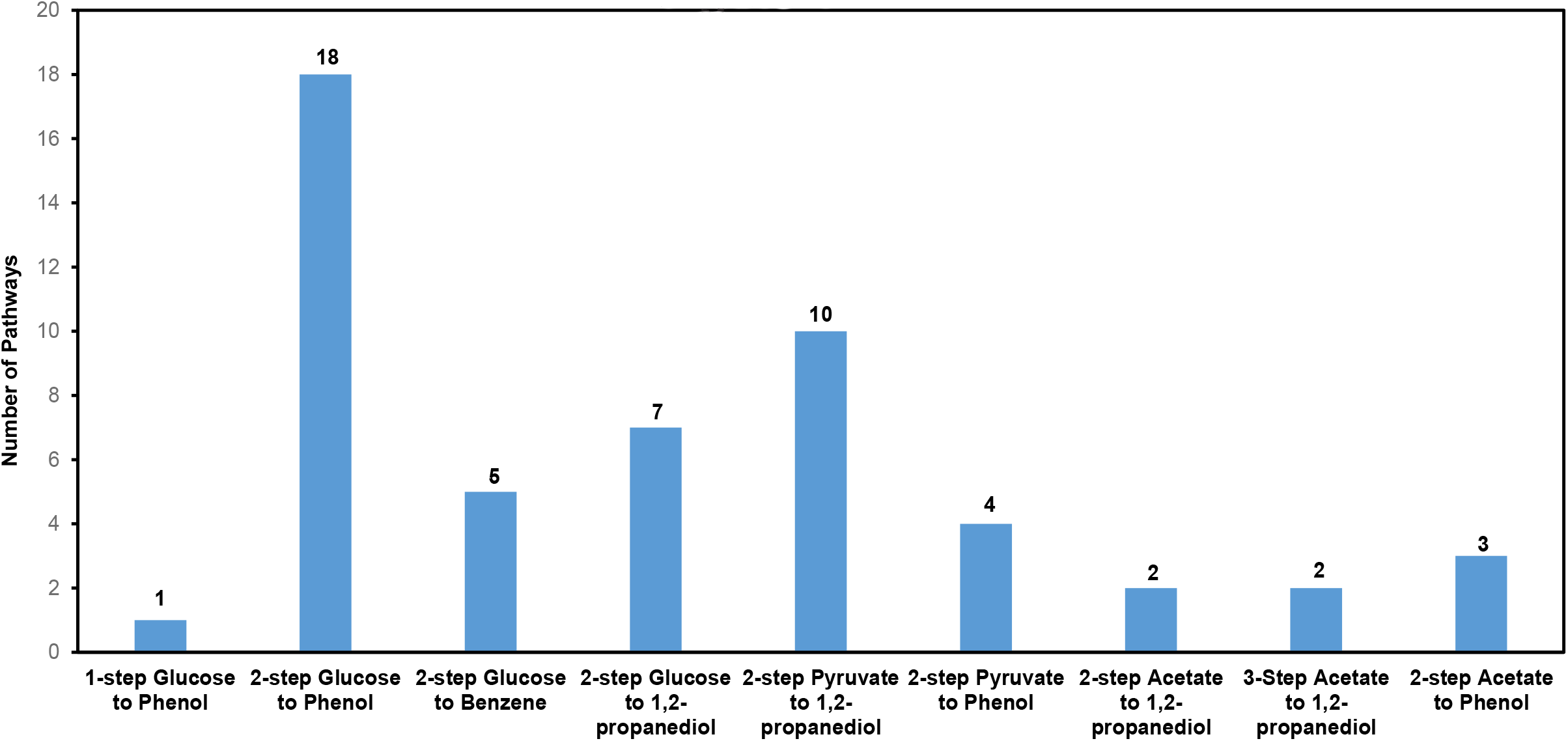
Number of accepted pathways to target commodity chemicals from glucose, pyruvate, and acetate. Different categories of accepted pathways to phenol, benzene, and 1,2-propanediol are shown from target starting compounds: glucose, pyruvate, and acetate. Each of these pathways are thermodynamically feasible and passed the pathway curation criteria.

The thermodynamic feasibility of the accepted pathways was analysed based on the overall standard Gibbs free energy of reactions 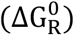 of each pathway. The standard Gibbs free energy of both the ReactPRED and RetroPath2.0-generated reactions were estimated using the group contribution method (Jankowski et al., 2008; Noor et al., 2012). Figure 6 shows a few notable examples of the accepted pathways discussed in this section, while Figure 7 depicts their thermodynamic feasibility information. Additional information of the pathways discussed in this section can be found in the supplementary data provided.

**Figure 6:**
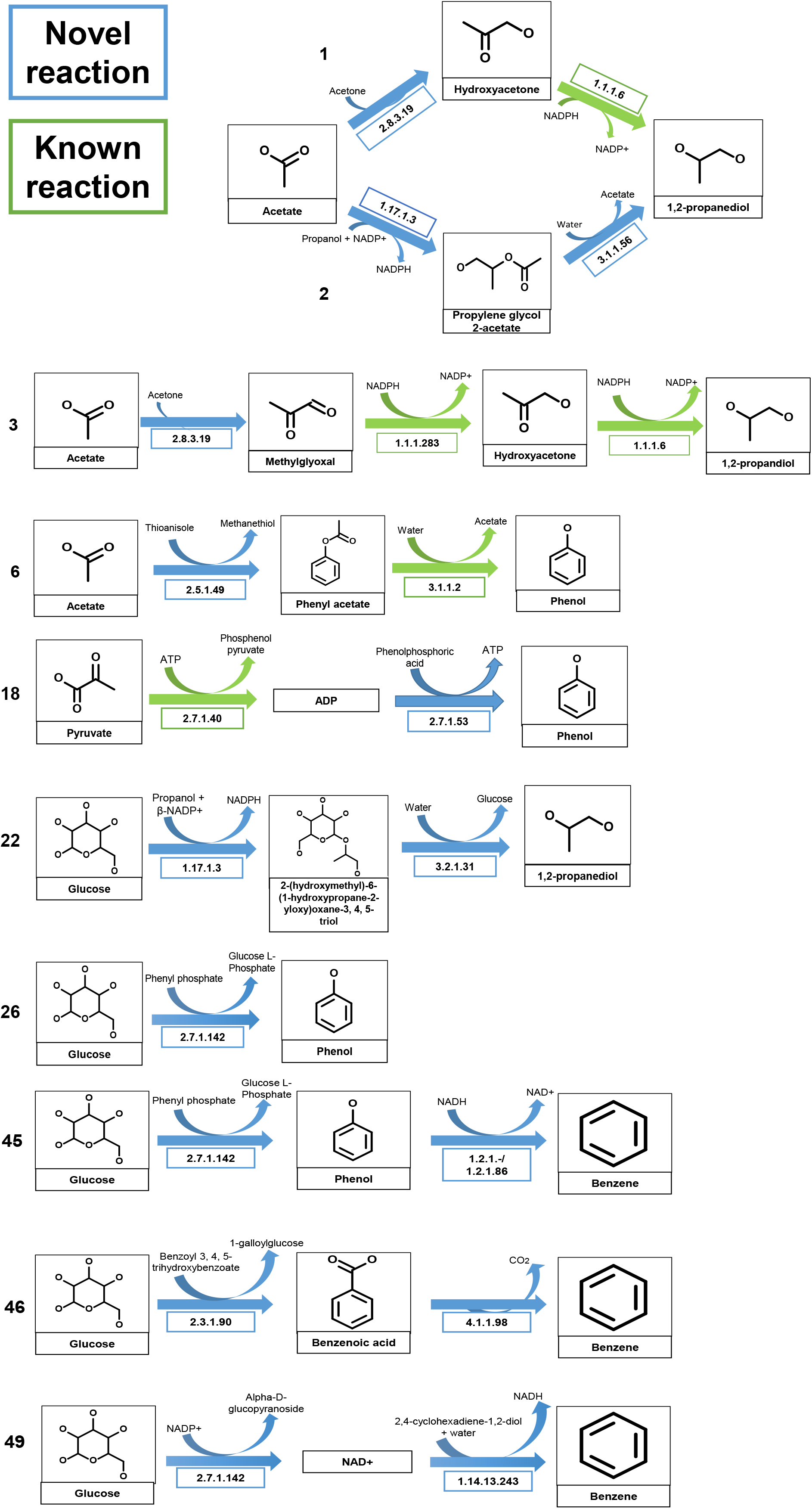
Examples of a few predicted pathways with assigned EC numbers. Each pathway contains at least one novel step (blue), while pathways 1, 3, 6, and 18 contain biologically known steps (green) as well. Pathways 22, 26, 45, 46, and 49 include only novel steps.

**Figure 7:**
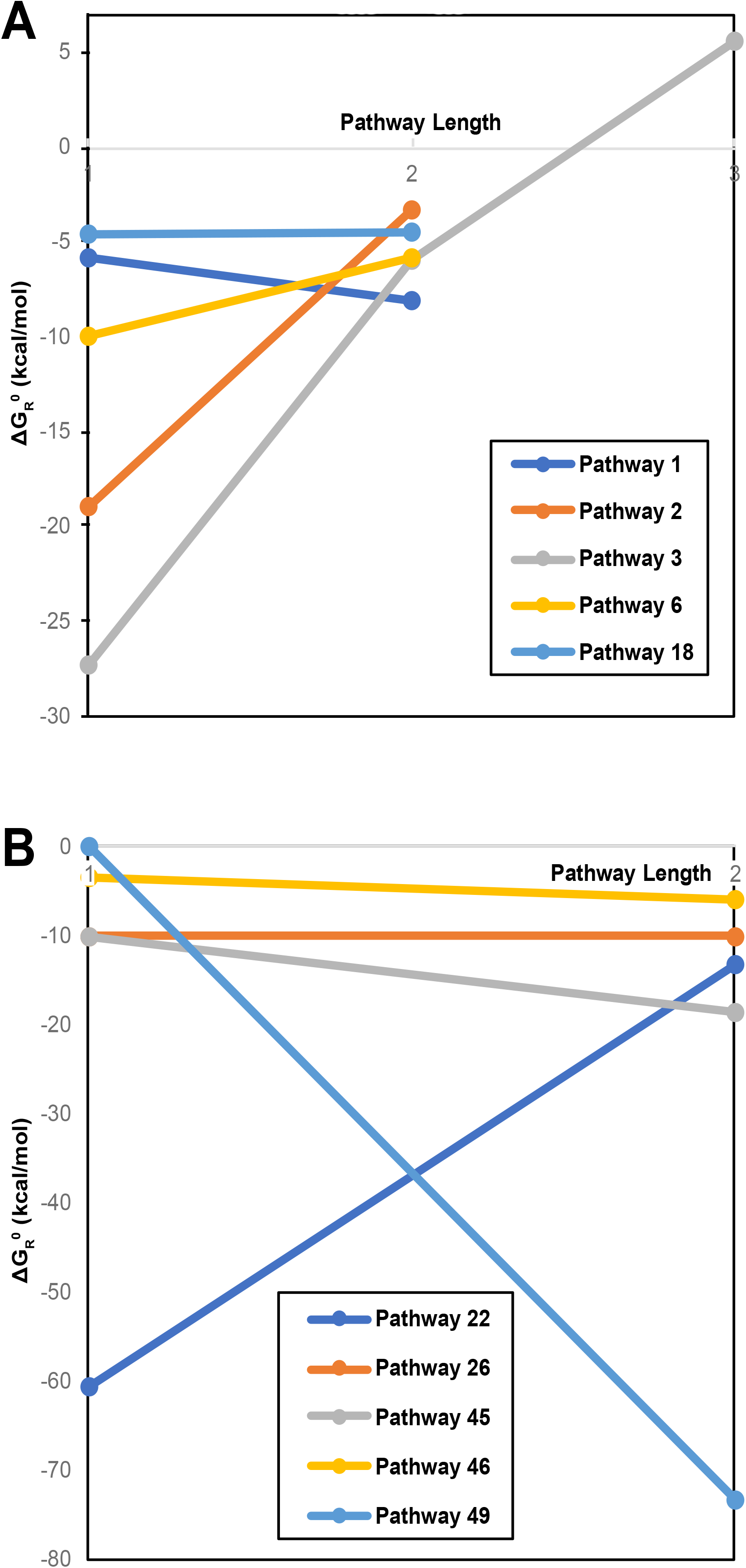
Analysis of the thermodynamic feasibility of the pathways shown in Figure 6. Pathways 1-3, 6, and 18 are shown in (A), and pathways 22, 26, 45-46 and 49 are shown in (B). The overall thermodynamic feasibility of each pathway was evaluated by estimating the standard Gibbs free energy of reaction 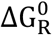 for each step followed by combining the 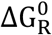 values for all relevant reactions in a pathway (see text for details).

Pathways 1-7 (Supplementary data) are the predicted pathways from acetate to 1,2-propanediol and phenol. Pathways 1, 3, and 4 are composed of the same novel reaction in the 1^st^ step, while pathway 2 include only novel reactions (Figure 6). Each acetate to 1, 2-propanediol producing pathway is thermodynamically feasible although pathways 3 and 4 include reaction steps with positive 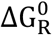 (Figure 7). Further comparison of pathways 1, 3, and 4 shows that the acetate to methylglyoxal reaction is thermodynamically more favourable than the acetate to hydroxyacetone reaction, resulting in a larger negative 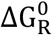 for the corresponding pathways. The first reaction step in pathways 5, 6, and 7 includes a novel reaction. This novel step involves the transferring of alkyl or aryl groups in 5 and 6, while in 7, this novel step uses a haloacetate dehalogenase-catalysed reaction. All three pathways generate phenol through an arylesterase reaction in the final reaction step. Each pathway is overall thermodynamically feasible. However, pathway 5 was found to be the most thermodynamically feasible while pathway 7 was the least of the 3 acetate to phenol pathways.

Pathways 8-21 (Supplementary data) are the predicted pyruvate pathways connecting pyruvate to the target commodity chemicals. Pathways 8-12 are predicted to produce hydroxyacetone in the first reaction step using the same novel reaction. Pathways 13-17 predict the production of lactic acid from pyruvate in the first reaction step. This step is a novel reaction for pathways 13-16, while it is a biologically known reaction in pathway 17. Only pathways 9 and 10 include a known reaction in the 2^nd^ reaction step. All pyruvate to 1, 2-propanediol pathways are overall thermodynamically feasible with each pathway having an overall negative 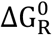. However, further comparison of the reaction steps revealed that the pathway producing 1, 2-propanediol via lactic acid were thermodynamically more favourable than the pathways through hydroxyacetone.

Pathways 18-21 produce phenol from pyruvate. The first reaction step in pathways 18, 21, and the 2^nd^ in pathway 20 are a known reaction, while the other reactions are novel in these pathways. Pathway 19 is predicted to use phenyl phosphono hydrogen phosphate as a co-reactant (Supplementary data). Phenyl phosphono hydrogen phosphate was only identified in the PubChem database, indicating that it is not a natural biological compound. Examining the thermodynamic feasibility of the pyruvate to phenol pathways reveal that pathways 18, 19, and 20 have the same 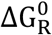 (−4.61 and −4.5 kcal/mol) for the first and second reaction steps, while only pathway 21 has different 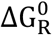 values (−4.6 and −1.7 kcal/mol) for the two reactions. These estimates lead to the fact that pathways 18, 19, and 20 are overall thermodynamically more favourable than pathway 21.

All predicted glucose to 1, 2-propanediol pathways, i.e., pathways 22-25 were retrosynthetically generated using ReactPRED and are composed of two novel reactions in both steps. Notably, all of these pathways contain a compound in the first reaction step only found in the PubChem database: 2-(hydroxymethyl)-6-(1-hydroxypropane-2-yloxy)oxane-3,4,5-triol (pathway 22 in Figure 6). The reactions in pathways 22-25 are all thermodynamically feasible with pathway 22 is estimated to be the most thermodynamically feasible glucose to 1, 2-propanediol pathway while pathway 25 is the least. The presence of NADP^+^ as a cofactor in the first reaction step of pathway 22 is likely to make it thermodynamically more favourable, as NADP^+^ works alongside enzymes to provide energy for cellular reactions (Xiao et al., 2018).

The only 1-step pathway from glucose to phenol (Pathway 26 in Figure 6) was generated retrosynthetically using ReactPRED. Pathways 27-32 include a known reaction in the first step, while the second reaction step in these pathways are all novel (Supplementary data). All of these pathways utilised water produced from the first step as a reactant for the second reaction. Pathways 33-40 and 44 are predicted to consist of novel reactions in both reaction steps. Interestingly, pathways 33-40 generated phenol from phenyl-α-D-glucoside in the second step using the same reaction; this reaction was classified as α-galactosidase and was assigned EC 3.2.1.21. Pathway 41 and 42 included the same known reaction in the second reaction step, while this 2^nd^ step in pathway 43 was identified as reaction R05626 using the KEGG database. Overall, each glucose to phenol pathway is thermodynamically feasible even though some pathways include reactions with a positive 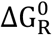.

The second step in pathways 45 and 46 were retrosynthetically generated by the RetroPath2.0 software in the first iteration (Figure 6). These pathways were compared to the ReactPRED generated pathways to find similarities and construct novel reactions. Pathway 45 was confirmed by creating a customised reaction rule set and generating the reactions using ReactPRED’s pathway prediction system (Sivakumar et al., 2016), while pathway 46 was confirmed through comparison of ReactPRED’s retrosynthetic pathway results. Pathway 47-49 were all retrosynthetically generated by ReactPRED and consisted of two novel reactions. Each glucose to benzene pathway is overall thermodynamically feasible. Pathway 48 was identified as the most thermodynamically feasible glucose to benzene pathway with an estimated overall 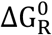 of −105 kcal/mol, while pathway 46 was the least thermodynamically feasible having an overall 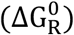 of −9.4 kcal/mol.

A closer examination of the accepted pathways further revealed similarities in the co-substrate use and bond transformation information, as well as in the estimated 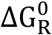 for multiple reaction steps. For example, 32 of the 80 accepted glucose pathways contain a reverse sucrose alpha-glucohydrolase reaction in the first reaction step to produce sucrose and water (Supplementary data). These pathways used water as the starting compound in the next reaction step and compounds only identified in the PubChem database as co-reactants. Similarly, pathways 22-25, 34, and 46-49 included co-reactants that were only identified in the PubChem database. The presence of a compound only in PubChem but not in other biological databases (MetaCyc, MetaNetX, KEGG) implies that the compound is man-made or synthetic and may not be made by biological systems. 16 out of 28 glucose pathways contained one of these compounds, whilst none of the acetate or pyruvate pathways contained these compounds. Moreover, many other pathways, such as pathways 6-8, 23-26, 28-30 produce target compounds using commodity chemicals as co-reactants or generate CO_2_ (Pathway 46 in Figure 6) in their reaction steps. Thus, the proposed pathways, although novel, may not necessarily be considered ‘green’. Finally, most of the retrosynthetically generated pathways, including pathways 1, 13-16, 19, 22-26, 33-39, 40, 43-45 are completely composed of novel reactions, indicating that retrosynthetic generation allows for more potential novel reactions to be uncovered.

### Comparative analysis of ReactPRED and RetroPath2.0

Both software tools, ReactPRED and RetroPath2.0 were applied to generate novel biochemical pathways to three industrially important commodity chemicals: benzene, phenol and 1, 2-propanediol. These cheminformatics tools were designed to be user-friendly and customisable to conduct user-specific pathway design tasks (Delépine et al., 2018; Sivakumar et al., 2016). Additionally, both tools have unique features that enable the design of biochemical pathways of various lengths from different starting compounds to different targets.

Both ReactPRED and RetroPath 2.0 allow users to customise their inputs to generate reactions. For ReactPRED, the input reaction rules were completely customisable, and the software’s reaction rule creation system allows the user to generate tailored reaction rules for specific tasks (Sivakumar et al., 2016). However, this study only utilised ReactPRED’s default reaction rule set to generate outputs, i.e., compounds, reactions, and pathways. Additionally, ReactPRED’s default reaction rule set was created using reactions present in the MetaCyc database (Caspi et al., 2020), and identical rules were merged together; thus, indicating that the novel reactions may be catalysed by more than one enzyme. Uniquely, ReactPRED estimates the overall Gibbs free energy change of the generated reactions, allowing users to assess the thermodynamic feasibility of the generated outputs. Further, ReactPRED allows users to view and search for pathways based on thermodynamic feasibility, molecular weight, and substructure through the user-friendly pathway analysis system.

Comparatively, RetroPath2.0 allows the user to tailor not only the reaction rules but also the ‘sink’ compounds to find novel pathways in the context of a specific chassis organism. This study used the software’s default reaction rule set and sink compounds that were developed based on the genome-scale metabolic model of *E. coli*, iJO1366 (Orth et al., 2011) and MetaNetX (Moretti et al., 2016), a meta-database consisting of reactions extracted from the KEGG, MetaCyc, Rhea (Lombardot et al., 2019) and Reactome (Jassal et al., 2020) databases. Additionally, RetroPath2.0 automatically assigns an EC number to each reaction within the chassis strain. Uniquely, RetroPath2.0 uses ‘sinks’ to prevent further iterations of the algorithm using compounds found within the chassis strain. This strategy, thus, not only shortens the execution time of the algorithm, but also prevents the combinational explosion of pathways that is usually generated with cheminformatics tools.

Although ReactPRED and RetroPath2.0 have benefits to aid biochemical pathway design, both software tools have limitations. For example, both pieces of software generate compounds that do not exist in nature, i.e., the compounds are unidentifiable in both biochemical and chemical databases. This is an important limitation, as these compounds and relevant reactions cannot be removed automatically from the generated results. Therefore, each pathway is required to be individually analysed to find and discard the reactions and pathways containing these compounds. Another major limitation of ReactPRED is its longer execution time, which can take anywhere from a few seconds to a week to generate the desired outputs. The time taken for ReactPRED to generate outputs is dependent on the number of potential bond transformations available for the starting compound and the number of reaction rules used to predict reactions. For instance, larger input molecules with more bond transformation potential will take longer to predict reactions. Furthermore, similar to many other cheminformatics tools, ReactPRED suffers from the combinational explosion of predicted reactions and pathways. As the pathway length increases, the number of predictions could be in the millions, leading to a greater effort to sift through the data to find meaningful results. Additionally, assigning EC numbers to the ReactPRED predicted reactions is also challenging and requires extensive analysis of the reactions, as discussed in this study, to assign a complete EC number.

A significant limitation of both pieces of software is that neither ReactPRED nor RetroPath2.0 can propose targeted pathways, i.e., generate only the desired reactions connecting starting compounds to target compounds automatically. Instead, both algorithms will continue to generate reactions iteratively until they are converged based on specific cut-off parameters such as pathway length and bond transformation diameter. Hence, a substantial amount of downstream pathway curation work is required to find the meaningful and novel results.

## Conclusions

Cheminformatics tools, ReactPRED and RetroPath2.0 were utilised to design novel biochemical pathways to produce three industrially important commodity chemicals with limited biochemical knowledge: benzene, phenol, and 1, 2-propanediol. All of the 49 designed pathways from glucose, acetate, and pyruvate contained at least one novel step, i.e., biologically unknown reaction, and all were found to be thermodynamically feasible. A novel methodology for curation of thousands and millions of pathways generated by both software tools was developed, and this method can be used as a guide for designing biochemical pathways to produce not only commodity chemicals but also nutraceuticals and pharmaceuticals. RetroPath2.0 and ReactPRED were also comparatively assessed to provide further insight on their effectiveness as a biochemical pathway design tool, as well as their advantages and limitations in the context of a specific design task. Although both software tools are user-friendly and help design novel pathways, these tools also produce thousands of pathways with compounds non-existent in nature. Hence, this study can be used to develop practical pathway curation strategies while using similar cheminformatics tools to design biochemical pathways. Moreover, the designed pathways can be used as valuable hypotheses for experimental implementation of the pathways in suitable chassis organisms for sustainable production of bio-based commodity chemicals.

## Notes

### Competing Interest Statement

The authors have declared no competing interest.

